# Discovery of influenza drug resistance mutations and host therapeutic targets using a human airway chip

**DOI:** 10.1101/685552

**Authors:** Longlong Si, Rachelle Prantil-Baun, Kambez H Benam, Haiqing Bai, Melissa Rodas, Morgan Burt, Donald E. Ingber

## Abstract

Here we demonstrate that influenza virus replication, host responses to infection, evolution through mutation or gene reassortment, and clinical efficacy of antiviral drugs can be reconstituted in a human Airway Chip microfluidic culture device. Modeling human-to-human transmission of infection in the continued presence of antiviral drugs on chips led to the emergence of clinically prevalent mutations responsible for amantadine- and oseltamivir-resistance, as well as the discovery of new resistance mutations. Analysis of infection responses resulted in identification of host therapeutic targets and demonstration that existing non-antiviral drugs may be repurposed to inhibit viral replication and synergize with antiviral therapeutics by targeting the host response to infection rather than the virus itself. This Influenza Chip may represent an alternative preclinical tool for development of new antiviral drugs and vaccines.

**One Sentence Summary:** New drug resistance mutations and potential tolerance-inducing therapeutics were discovered using an organ chip model of influenza infection.

## Main Text

The greatest challenge for combating influenza virus infection is rapid virus evolution among human populations, which leads to the emergence of mutated viruses making existing anti-influenza drugs and vaccines ineffective (*1, 2*). Development of more effective means to control influenza will therefore require better prediction of virus evolution of resistance to therapy and more rapid development of novel drugs and vaccines, both of which are currently limited by the lack of clinically relevant preclinical models (*3*). Here we explored whether human Organ-on-a-Chip (Organ Chip) microfluidic culture technology (*4-7*) can be used to develop a physiologically and clinically relevant *in vitro* model of human influenza A infection for prediction of influenza evolution and preclinical evaluation of anti-influenza therapeutics.

Using a previous protocol with some modification (*5*), we first built a human lung Airway Chip containing 2 parallel microchannels separated by an extracellular matrix (ECM)-coated membrane lined by primary human lung airway epithelial cells (HLAECs) cultured under an air-liquid interface (ALI) on one side within the ‘airway’ channel, with human pulmonary microvascular endothelial cells (HPMVECs) grown on the other in the presence of continuous medium flow to mimic vascular perfusion with or without human immune cells within the ‘vascular’ channel (**Fig. 1A**). After 3 weeks of culture, the HLAECs differentiated into a mucociliary, pseudostratified epithelium with cells linked by continuous ZO1-containing tight junctions (**fig. S1A**) and proportions of airway-specific cell types similar to what is observed in vivo (**fig. S1B**), and the underlying endothelium formed a planar cell monolayer joined by VE-containing adherens junctions (**fig. S1A**). Differentiation of the airway epithelium on-chip resulted in development of a high level of barrier function (**fig. S1C**) and mucus secretion (**fig. S1D**), as well as elevated expression of multiple serine proteases that are essential for the activation and propagation of influenza viruses *in vivo* (*8-10*), including TMPRSS2, TMPRSS4, TMPRSS11D and TMPRSS11E (DESC1), when compared to either undifferentiated HLAECs or MDCK cells (**fig. S1E**). Under these microfluidic culture conditions, these highly differentiated human airway structures and functions can be maintained for more than 2 months *in vitro*.

**Fig. 1.**
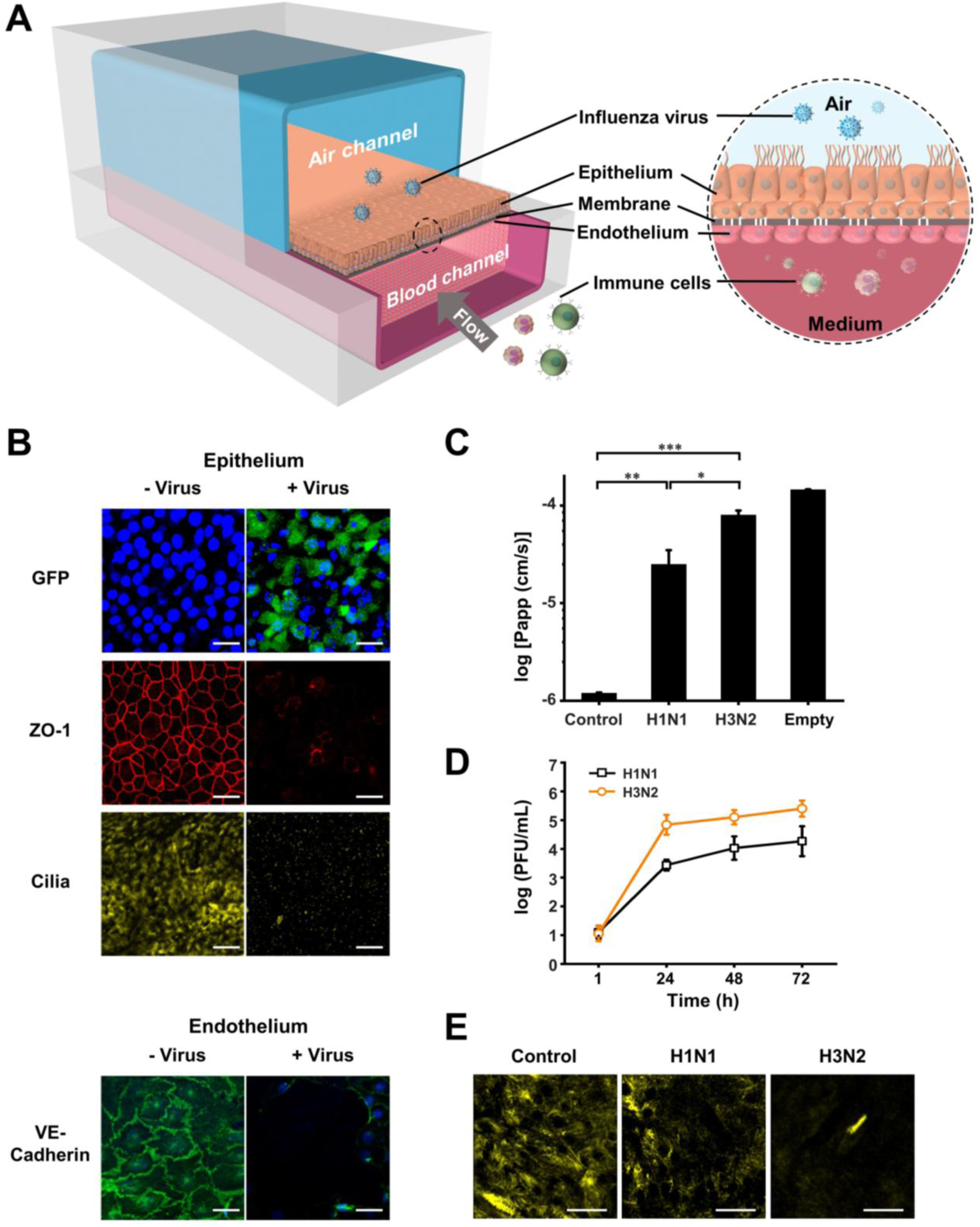
Influenza infection in the human airway chip. (**A**) Schematic diagram of a cross-section through the airway chip. (**B**) Immunofluorescence micrographs showing the effects of infection with GFP-labeled influenza H1N1 virus (MOI = 0.1) on distribution of ZO1-containing tight junctions and cilia in epithelium and VE-cadherin-containing adherens junctions in endothelium of the human airway chip at 48 h post-infection. (**C**) Increase in barrier apparent permeability (log P_app_) within the human Airway Chip measured 48 h post-infection with H1N1 or H3N2 virus (MOI = 0.1) compared to no infection (Control) or a chip without cells (Empty). (**D**) Replication kinetics of influenza A/WSN/33 (H1N1) and A/Hong Kong/8/68/ (H3N2) virus (MOI = 0.001) in human Airway Chips. (**E**) Immunofluorescence micrographs showing apical cilia 24 h post-infection with H1N1 or H3N2 (MOI = 0.1). Bar, 50 µm; *, P<0.05; **, P<0.01; ***, P<0.001.

We then inoculated the airway epithelium with GFP-labeled influenza H1N1 (PR8) virus (*11*) through the upper air channel of the microfluidic chip to mimic *in vivo* infection with airborne influenza (**Fig. 1A**). Real-time fluorescence microscopic analysis confirmed viral infection of the human airway epithelium as indicated by GFP expression (**Fig. 1B and movie S1**), and this was accompanied by damage to the epithelium, including disruption of tight junctions, loss of apical cilia, and compromised barrier function (**Fig. 1B,C**). Influenza infection also led to disruption of the endothelium as evidenced by loss of VE-cadherin containing adherens junctions (**Fig. 1B**), which was consistent with vascular leakage that is induced by influenza infections *in vivo* (*12, 13*). Analysis on replication kinetics of H1N1 and H3N2 viruses indicated that both viruses exhibited large (10,000-to 100,000-fold) increases in viral titers over 24 to 48 hours; however, H3N2 exhibited ∼10-fold greater replication efficiency (**Fig. 1D**) and caused more barrier function damage and cilia loss (**Fig. 1C,E**), which is consistent with the clinical observations that H3N2 is more infectious and causes more severe clinical symptoms in humans (*14*). Assessment of innate immune responses to infection of H1N1, H3N2 and H5N1 revealed that H3N2 and H5N1 viruses that produce more severe clinical symptoms than H1N1 induced higher levels of cytokines and chemokines, and the most virulent H5N1 induced the highest concentrations (**fig. S2**). These results mirror the clinical finding that patients infected with H5N1 have increased serum concentrations of these inflammatory factors relative to those with H1N1 or H3N2, which significantly contributes to disease pathogenesis (*14*). Importantly, donor-to-donor variability was minimal as similar results were obtained among chips created with HLAECs obtained from 5 different donors.

We next tested the potential of our human Airway Chip influenza model to enable the *in vitro* prediction of influenza virus evolution by mutation when viruses spread from patient-to-patient through human populations. Human patient-to-patient transmission of influenza virus was mimicked by passaging virus from chip to chip under the selection pressure of antiviral drugs (**Fig. 2A**). When we treated human airway chips infected with influenza A/WSN/33 (H1N1) virus with the clinically used anti-influenza drug, amantadine, at a dose (1 µM) that inhibits replication of this influenza strain by ∼90%, its inhibition rate decreased to ∼10% after 8 human chip-to-chip passages (**Fig. 2A,B**), indicating the emergence of a pool of amantadine-resistant viruses. Sequencing of these drug-resistant viruses led to the identification of 3 mutated virus strains with mutation sites located within the viral M2 protein (**Fig. 2C**), which is the known target of amantadine (*15*). Among them, a single mutation at S31N conferred amantadine-resistance with the half inhibitory concentration (IC_50_) increasing over 500-fold (from 47 nM to 24.7 µM) (**Fig. 2D**). Importantly, this is the same site mutation that has been frequently detected in clinically validated amantadine-resistant influenza viruses (*16*). In addition, we identified two previously unknown double mutants containing the S31N mutation, as well as either G34E or L46P substitutions (**Fig. 2C**). Interestingly, these mutations conferred even greater resistance to amantadine with the IC_50_ increasing over 1000-fold (from 47 nM to > 100 or 65 µM, respectively) (**Fig. 2D**), indicating that extended exposure to amantadine may induce the emergence of more highly resistant virus strains.

**Fig. 2.**
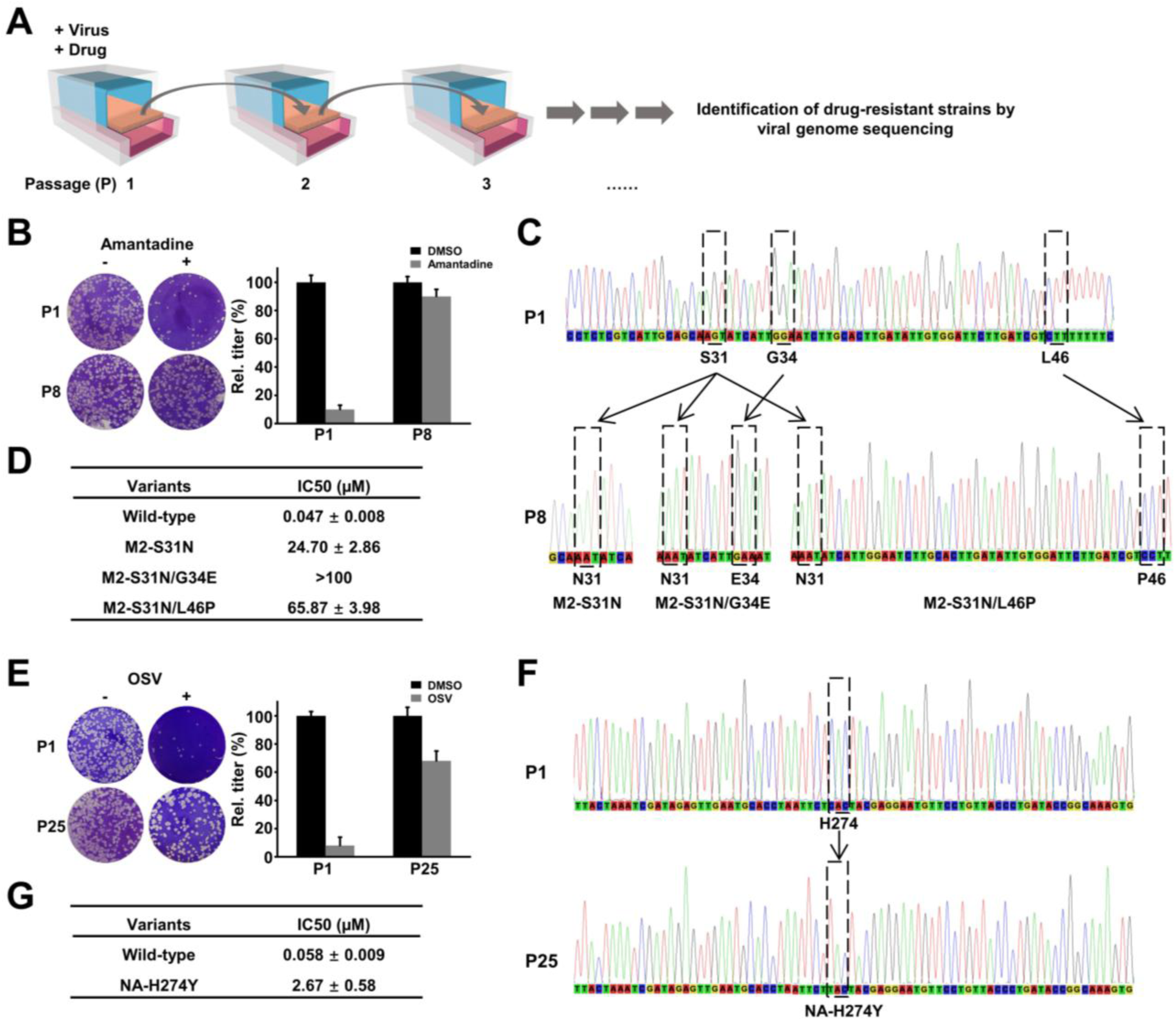
Modeling influenza virus evolution through mutation on-chip. (**A**) Schematic diagram of method used to generate and identify drug-resistant viruses by human chip-to-chip transmission under drug pressure. (**B**) Photographs (left) of plaques and graph (right) showing plaque titers of progeny virus at 1st and 8th passage (P) in control (-) versus amantadine-treated (+) chips. (**C**) Sequencing graphs showing three mutants(M2-S31N, M2-S31N/G34E, and M2-S31N/L46P) detected in the amantadine-resistant virus pool. (**D**) IC_50_ values of amantadine against parental strain and the three mutants. (**E**) Photographs (left) of plaques and graph (right) showing plaque titers of progeny virus at 1st and 25th passage in control (-) versus oseltamivir (OSV)-treated (+) chips. (**F**) Sequencing graphs showing one mutant (NA-H274Y) detected in the OSV-resistant virus pool. (**G**), IC_50_ values of OSV against parental strain and mutant.

We used the same approach to explore the propensity of the widely used anti-influenza drug, oseltamivir, to induce viral resistance, and found that the inhibition rate also decreased significantly (from ∼90% to ∼30%) over time (**Fig. 2E**). But this was a much slower process as 25 human chip-to-chip passages were required for resistance to develop (**Fig. 2E**). Sequencing of the oseltamivir-resistant virus pool revealed one strain with a mutation site located within the influenza viral neuraminidase (NA) protein that is the known target of oseltamivir (**Fig. 2F**) (*17*). The H274Y mutation conferred oseltamivir resistance with the IC_50_ increasing from 58 nM to 2.67 µM (**Fig. 2G**), and importantly again, the same mutation has been detected in clinical cases (*17*).

To explore whether the Airway Chip also could support influenza virus evolution through gene reassortment as occurs when different virus strains co-infect the same host (*18*), chips were co-infected with H3N2 and H1N1 viruses and their progeny viruses were then isolated and sequenced (**Fig. 3A**). Genotype analysis of 100 progeny viruses revealed that the H3N2 and H1N1 parental strains were predominant; however, 19 virus reassortants emerged that represented three distinct genotypes (3, 4, and 5) (**Fig. 3B**). These included seven H1N2 reassortants containing the hemagglutinin (HA) gene segment from H1N1 with the other gene segments from H3N2 in genotype 3; two new H3N2 reassortants containing the polymerase basic protein-1 (PB1) gene segment from H1N1 with others from H3N2 in genotype 4; and ten new H3N2 reassortants containing the matrix protein (M) gene segment from H1N1 with others from H3N2 in genotype 5 (**Fig. 3B**). These reassortants did not exhibit higher replication competence than their parental strains (**Fig. 3C**), but the emergence of antigenically drifted H1N2 reassortants led to complete resistance to H3N2-specific anti-HA antibody (**Fig. 3D**), indicating the potential risk of pandemic development if antigen-mismatched vaccination occurred. Importantly, the H1N2 reassortant predicted on-chip also has been detected in patients during influenza season in which H1N1 and H3N2 co-circulated (*18*). Even more concerning, a high frequency of double resistant viruses emerged that are resistant to both oseltamivir and amantadine as a result of reassortment of oseltamivir-resistant virus and amantadine-resistant virus when they were used to co-infect human airway chips (**fig. S3**). Thus, *in vivo* virus evolution through reassortment can be easily studied in this *in vitro* human influenza infection model.

**Fig. 3.**
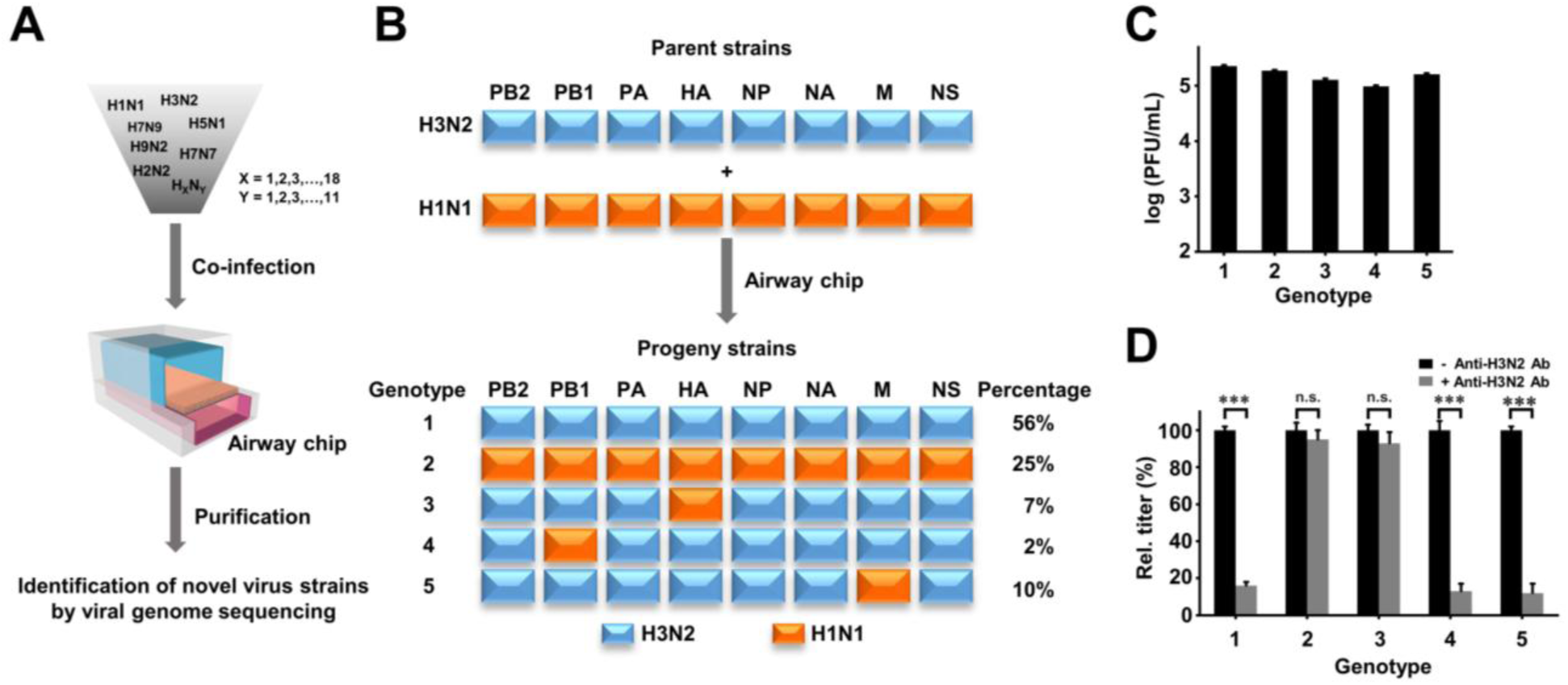
Modeling influenza virus evolution through gene reassortment on-chip. (**A**) Schematic diagram of the method for generation and identification of reassortants in Airway Chips co-infected by different strains. (**B**) Five genotypes with different percentages of incidence revealed from sequencing analysis of 100 progeny viruses isolated from Airway Chips co-infected by H1N1 and H3N2 (blue boxes, segments derived from H3N2; orange boxes, segments from H1N1). (**C**) Replication titers of different genotypes of reassortants and their parental strains in Airway Chips 48 h post-infection (MOI = 0.1). (D) The neutralization activity of anti-H3N2 HA antibody (10 µg/mL) against different genotypes of reassortants and their parental strains. ***, P<0.001; n.s., not significant.

Next, we tested the reliability of our model to evaluate anti-influenza therapeutics using oseltamivir and anti-HA antibody (**fig. S4**). As oseltamivir is normally metabolized to release its active metabolite, oseltamivir acid, by the liver *in vivo*, we introduced oseltamivir acid into the vascular channel of a human airway chip infected with H1N1 virus, mimicking its oral administration and distribution through the vasculature in flu patients. Oseltamivir efficiently inhibited viral replication (**fig. S4A**), protected barrier function of airway (**fig. S4B**), and maintained epithelial tight junctions (**fig. S4C**) in chips infected with influenza. In addition, an anti-H1N1 HA antibody significantly inhibited virus propagation, and it exhibited more potency against H1N1 than H3N2, as expected due to differences in antigen epitopes (**fig. S4D**).

The Airway Chip was then used to explore whether host serine proteases could serve as alternative targets for therapeutic intervention given their significantly elevated expression in human airway (**fig. S1E**), and their critical roles in influenza virus propagation (*8, 10*). We screened a library of serine protease inhibitors, and found that aprotinin/Trasylol, leupeptin, AEBSF and the clinically used anticoagulant, nafamostat exhibited anti-influenza efficacies against H1N1 and H3N2 on-chip (**Fig. 4A and fig. S5**). Nafamostat protected airway barrier function (**Fig. 4B**) and tight junction integrity (**Fig. 4C**), and decreased production of cytokines and chemokines (**Fig. 4D**), when the Airway Chips were challenged with influenza. Analysis of nafamostat’s mechanism of action revealed that it efficiently blocks the cleavage of influenza viral HA0 protein into HA1 and HA2 subunits mediated by serine proteases, including TMPRSS11D and TMPRSS2 (**Fig. 4E and fig. S6A to C**), a process that is essential for influenza virus infection (*19, 20*). This mechanism was also shared by the other protease inhibitors (**fig. S6D**).

**Fig. 4.**
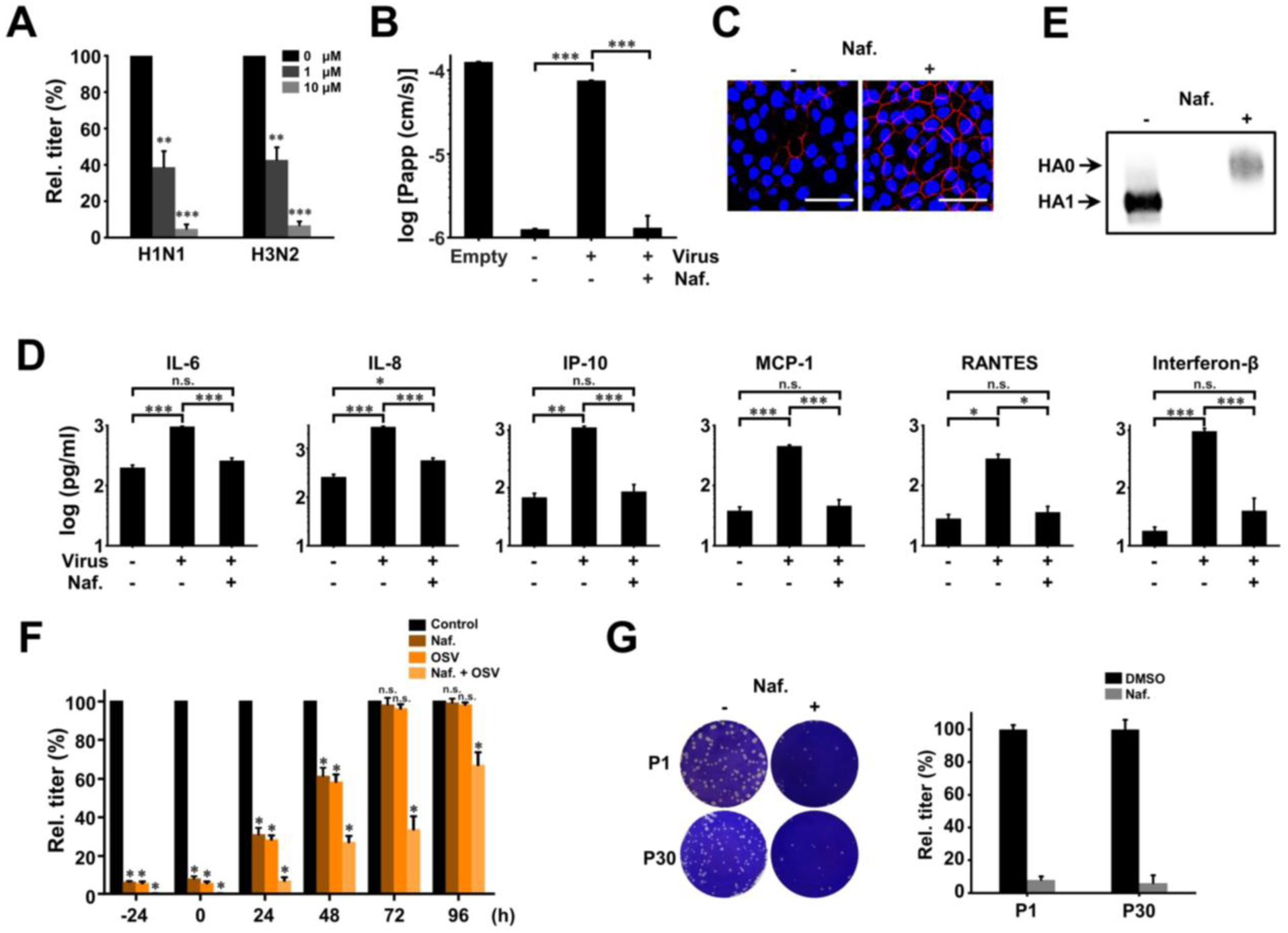
Repurposing of MMP inhibitors and nafamostat as anti-influenza therapeutics in the human Airway Chip. (**A**) Virus titer detection showing the effects of nafamostat on virus replication of H1N1 and H3N2 at 48 hours. (**B**) Barrier permeability (log P_app_) within the human Airway Chip measured 48 h post-infection with H1N1 (MOI = 0.1) in the presence/absence of 10 µM nafamostat versus in an empty chip. (**C**) Immunofluorescence micrographs showing preservation of ZO1-containing tight junctions in airway epithelium infected with H1N1 (MOI = 0.1) 48 h post-infection by treatment with nafamostat (10 µM). (**D**) Production of various influenza-associated cytokines and chemokines in the Airway Chip in the presence/absence of 10 µM nafamostat. (**E**) Western blot showing inhibition of cleavage of influenza HA0 into HA1 and HA2 subunits by nafamostat. (**G**) The effects on relative viral titers of nafamostat, oseltamivir and their combination when added to H1N1 virus-infected Airway Chips at indicated times; note the synergistic effects of these two drugs at later times. (**G**) Photographs (left) of plaques and graph (right) showing plaque titers of progeny virus at 1st and 30th passage in control (-) and nafamostat-treated (+) chips, and the lack of development of drug resistance. Bar, 50 µm. *, P<0.05; **, P<0.01; ***, P<0.001; n.s., not significant.

Oseltamivir is usually recommended for clinical use within 2 days of influenza infection (*21, 22*), and when we added this drug at different time points of influenza infection on-chip, it similarly was only effective when administered within 48 h post-infection (**Fig. 4F**). Nafamostat exhibited a similar 48 h therapeutic window; however, when these two drugs were combined, they exerted a more potent anti-influenza effect (**Fig. 4F**). Most importantly, by combining nafamostat with oseltamivir, we were able to extend the therapeutic window from 48 h to 96 h (**Fig. 4F**). Furthermore, no viral resistance to nafamostat was detected even when virus infection was transmitted over 30 chip-to-chip passages in the presence of this drug (**Figs. 2A and 4G**), which is ascribed to the highly conserved nature of host targets. This is significantly different from all current clinically used viral protein-targeting antiviral drugs, such as amantadine and oseltamivir, which induce rapid emergence of drug-resistant virus strains (**Fig. 2**) (*16, 17*).

Our results show that the human Airway Chip can be used to create a physiologically and clinically relevant *in vitro* model of human influenza infection. This experimental platform could be used to predict potentially emerging viruses that become resistant to current anti-influenza drugs or vaccines, and hence, cause influenza pandemics. By combining the influenza evolution prediction in Airway Chips with existing influenza surveillance approaches (*18, 23-25*), it might be possible to establish public databases of influenza evolution in humans, which could lead to better selection of candidate viruses for vaccines and early detection of drug-resistant viruses (*26*). This model also may enable preclinical evaluation of new anti-influenza therapeutics, leading to the identification of candidate anti-influenza drugs. It is important to note that the protease inhibitor drugs we identified to have antiviral activity have a novel mechanism of action in that they target the host, rather than the virus itself, and thus, increase host tolerance to infection. As some of these existing drugs have been approved by the FDA for other medical uses (*27*), they could be repurposed and moved rapidly to clinical testing as anti-influenza therapeutics either alone or in combination with existing antiviral agents.

## References and Notes

## Acknowledgments

We thank P. Palese and A. Garcia-Sastre for kindly sharing the influenza virus strains, and R.A.M. Fouchier and G.F. Gao for providing the influenza virus rescue systems.

## Author contributions

L.S., R.P., and D.E.I. designed and analyzed experiments; L.S. performed experiments and data analysis; K.H.B. identified the anti-influenza efficacy of nafamostat; H. B. analyzed the serine proteases; M.R. and M.B. validated the airway cells from different donors; L.S. and D.E.I. wrote the manuscript, with all authors providing feedback.

## Data and materials availability

Sharing of materials will be subject to standard material transfer agreements. The nucleotide sequences used in the study have been deposited in GeneBank under accession numbers CY034139.1, CY0334138.1, X17336.1, HE802059.1, CY034135.1, CY034134.1, D10598.1, M12597.1, CY176949.1, CY176948.1, CY176947.1, CY176942.1, CY176945.1, CY176944.1, CY176943.1, CY176946.1, DQ487334.1, DQ487333.1, DQ487335.1, DQ487340.1, DQ487339.1, DQ487337.1, and DQ487338.1, DQ487336.1. Additional data are presented in the Supplementary Materials.

## Supplementary Materials

Materials and Methods

Figures S1-S6

Tables S1-S2

Movies S1

## References

1. X. Du, A. A. King, R. J. Woods, M. Pascual, Sci Transl Med 9, (2017).

2. S. P. Layne, A. S. Monto, J. K. Taubenberger, Science 323, 1560–1561 (2009).

3. F. Krammer et al., Nat Rev Dis Primers 4, 3 (2018).

4. D. Huh et al., Science 328, 1662–1668 (2010).

5. K. H. Benam et al., Nat Methods 13, 151–157 (2016).

6. D. Huh et al., Sci Transl Med 4, 159ra147 (2012).

7. K. H. Benam et al., Cell Syst 3, 456–466 e454 (2016).

8. M. Dittmann et al., Cell 160, 631–643 (2015).

9. J. Zhou et al., Proc Natl Acad Sci U S A 115, 6822–6827 (2018).

10. R. W. Chan, M. C. Chan, J. M. Nicholls, J. S. Malik Peiris, Virus Res 178, 133–145 (2013).

11. B. Manicassamy et al., Proc Natl Acad Sci U S A 107, 11531–11536 (2010).

12. S. M. Armstrong, S. Mubareka, W. L. Lee, Antiviral Res 99, 113–118 (2013).

13. W. Zhu et al., Nature 492, 252–255 (2012).

14. C. Y. Cheung et al., Lancet 360, 1831–1837 (2002).

15. S. D. Cady et al., Nature 463, 689–692 (2010).

16. R. A. Bright et al., Lancet 366, 1175–1181 (2005).

17. P. J. Collins et al., Nature 453, 1258–1261 (2008).

18. G. Neumann, T. Noda, Y. Kawaoka, Nature 459, 931–939 (2009).

19. T. M. Tumpey et al., Science 310, 77–80 (2005).

20. D. C. Ekiert et al., Science 333, 843–850 (2011).

21. J. H. Beigel et al., Lancet Infect Dis 17, 1255–1265 (2017).

22. W. Simonson, Geriatr Nurs 40, 99–100 (2019).

23. M. Luksza, M. Lassig, Nature 507, 57–61 (2014).

24. D. J. Smith et al., Science 305, 371–376 (2004).

25. J. C. Obenauer et al., Science 311, 1576–1580 (2006).

26. S. Yamayoshi, Y. Kawaoka, Nat Med 25, 212–220 (2019).

27. F. A. DeLano, D. B. Hoyt, G. W. Schmid-Schonbein, Sci Transl Med 5, 169ra111 (2013).

28. H. J. Kim, D. Huh, G. Hamilton, D. E. Ingber, Lab Chip 12, 2165–2174 (2012).

29. S. Chutinimitkul et al., J Virol 84, 11802–11813 (2010).

30. L. Si et al., Science 354, 1170–1173 (2016).

