## Supplementary file for "Discovery of influenza drug resistance mutations and host therapeutic targets using a human airway chip"

**This PDF file includes:**

Materials and Methods

Figs. S1 to S6

Tables S1 to S2

Captions for Movies S1

**Other Supplementary Materials for this manuscript include the following:**

Movies S1

### Materials and Methods

#### Human Airway Chip Fabrication

Microfluidic Organ Chip devices were fabricated using poly-dimethyl siloxane (PDMS; Sylgard 184 silicone elastomer kit) as previously described using soft lithography (*4*,*5*) or obtained from Emulate Inc. (Boston, MA). The device contains two adjacent parallel microchannels (apical, 1 mm wide × 1 mm high; basal, 1 mm wide × 0.2 mm high; length of overlapping channels, 16.7 mm). The two channels are either separated by a 50 µm thick porous PET membrane (0.4 µm holes; $4 \times 10^6$ pores/cm$^2$) or PDMS membrane (7 µm holes with 40 µm spacing). Both channels were washed with 70% ethanol, filled with 0.5 mg/mL ER1 solution (Emulate Inc.) in ER2 buffer (Emulate Inc.) and placed under UV lamp (Nailstar, NS-01-US) for 20 min to activate the surface. The channels were then washed sequentially with ER2 buffer and PBS. The porous membranes were coated on both sides with collagen type IV from human placenta (0.5 mg/mL; Sigma-Aldrich) at room temperature for overnight. The solution was then aspirated from the chip and the chip was ready for seeding cells.

#### Cell culture

Primary HLAECs (Lonza, USA) obtained from healthy donors 448571, 446317, 623950, 485960, and 672447) or a COPD donor (370751) were expanded in 75-cm$^2$ tissue culture flasks using airway epithelial cell growth medium (Promocell, Germany) until 60-70% confluent. Primary HPMVECs (Cell Biologics, USA) were expanded in 75-cm$^2$ tissue culture flasks using human endothelial cell growth medium (Cell Biologics, USA) until 70-80% confluent. To create the human Airway Chips, primary HPMVECs ($2 \times 10^7$ cells/mL) were first seeded in the bottom channel of the chip by inversion for 4 h in human endothelial cell growth medium, followed by seeding of the top channel with primary HLAECs ($2.5 \times 10^6$ cells/mL) for 4 h in airway epithelial cell growth medium. The respective medium for each channel was refreshed and the chips were incubated at 37℃ in 5% $CO_2$ under static condition for overnight. The attached cells were then continuously perfused with the respective cell culture medium for 5-7 days using an IPC-N series peristaltic pump (Ismatec) at a volumetric flow rate of 60 μL/h. Then the apical medium was removed and an ALI was established. The HLAECs were cultured at the ALI for 3-4 weeks with a constant flow of PneumaCult-ALI medium (StemCell) supplemented with 0.1% VEGF, 0.01% EGF, and 1mM $CaCl_2$ from an Endothelial Cell Medium Kit (Cell Biological, M1168) in the bottom channel. The chips were incubated in an incubator containing 5% $CO_2$ and 16-18% $O_2$ at 85-95% humidity, and the apical surface of the epithelium was rinsed once weekly with PBS to remove cellular debris and mucus.

**Immunofluorescence microscopy**

Cells were washed with PBS through the apical and basal channels, fixed with 4% paraformaldehyde (Alfa Aesar) for 20-25 min, and then washed with PBS before being stored at 4$^{o}$C until use. Fixed tissues were permeabilized on-chip with 0.1% Triton X-100 (Sigma-Aldrich) in PBS for 5 min, exposed to PBS with 10% goat serum (Life Technologies) and 0.1% Triton X-100 for 30 min at room temperature, and then incubated with corresponding primary antibodies (**table S1**) diluted in incubation buffer (PBS with 1% goat serum and 0.1% Triton X-100) overnight at 4$^{o}$C, followed by incubation with corresponding secondary antibodies (**table S1**) for 1 h at room temperature, and nuclei were counterstained with DAPI (Invitrogen) after secondary antibody staining. Fluorescence imaging was carried out using a confocal laser-scanning microscope (SP5 X MP DMI-6000, Germany) and image processing was done using Imaris (Bitplane, Switzerland).

**Mucus quantification**

Mucus present in the airway channel was isolated by infusing 50 µl PBS into the upper channel of the Airway Chip, incubating for 1 h at 37°C, and then collecting the fluid and storing it at -80°C before analysis. Quantification of mucus production was carried out by using Alcian Blue Staining (Thermo Fisher Scientific) and comparing to serially diluted standards of mucin (Sigma-Aldrich) in PBS (*28*).

**Barrier function assessment**

To measure barrier permeability, 50 µl cell medium containing Cascade blue (607 Da) (50 µg/mL; Invitrogen) was added to bottom channel and 50 µl cell medium was added to top channel. 2 h later, fluorescence intensity of medium of top and bottom channels was measured in three different small airway chips. The apparent permeability was calculated using the formula: $P_{app} = J/(A \times \Delta C)$, where $P_{app}$ is the apparent permeability, $J$ is the molecular flux, $A$ is the total area of diffusion, and $\Delta C$ is the average gradient.

**Quantitative reverse transcription-polymerase chain reaction (qRT-PCR)**

Total RNA was extracted from differentiated human Airway Chips, pre-differentiated HLAECs, or MDCK cells using TRIzol (Invitrogen). cDNA was then synthesized using AMV reverse transcriptase kit (Promega) with Oligo-dT primer. To detect cellular gene-expression level, quantitative real-time PCR was carried out according to the GoTaq qPCR Master Mix (Promega) with 20 µl of a reaction mixture containing gene-specific primers (**table S2**). PCR conditions were 1 cycle at 95℃ for 5 min, followed by 40 cycles at 95℃ for 15 s, 60℃ for 1 min, and 1 cycle at 95℃ for 15 s, 60℃ for 15 s, 95℃ for 15 s. Cellular mRNA expression levels were normalized with GAPDH levels.

**Virus propagation**

Influenza A virus strains used in this study include A/PR/8/34 (H1N1), GFP-labeled A/PR/8/34 (H1N1), A/WSN/33 (H1N1), A/Netherlands/602/2009 (H1N1), A/Hong Kong/8/68/ (H3N2), A/Panama/2007/99 (H3N2), and A/Hong Kong/156/1997 (H5N1). H3N2 viruses were obtained from the American Type Culture Collection (ATCC) and the Centers for Disease Control and Prevention (CDC). The other viruses were generated using reverse genetics techniques as previously described (11,*29*,*30*), and propagated in MDCK cells. When >90% cytopathic effect (CPE) was observed, the supernatants containing viruses were harvested, centrifuged at 1000 g for 10 min to remove cell debris, and the remaining viruses were stored at -80℃.

**Plaque formation assay**

Virus titers were determined by plaque formation assay. Confluent MDCK cell monolayers in 12-well plate were washed with PBS, inoculated with 1 mL of 10-fold serial dilutions of influenza virus samples, and incubated for 1 h at 37℃. After unabsorbed virus was removed, the cell monolayers were overlaid with 1 mL of DMEM (Gibco) supplemented with 1.5% low melting point agarose (Sigma-Aldrich) and 2 μg/mL TPCK-treated trypsin (Sigma-Aldrich). After incubation for 2-4 days at 37℃ under 5% $CO_2$, the cells were fixed with 4% paraformaldehyde, and stained with crystal violet (Sigma-Aldrich) to visualize the plaques; virus titers were determined as plaque-forming units per milliliter (PFU/mL).

**Influenza virus infection and treatment in the human Airway Chip**

Human Airway Chips were infected with influenza viruses by flowing 30 uL of PBS containing the indicated multiplicity of infection (MOI) of viral particles into the apical channel, incubating for 2 h at 37℃, and then removing the medium to reestablish an ALI. To measure virus propagation, the apical channel was incubated with 50 μl of PBS for 1 h at 37℃ at various times, and then both the apical fluid and vascular effluent were collected from the apical and basal channels, respectively, to quantify viral load using the plaque formation assay and analyze released cytokines and chemokines. The tissues cultured on-chip were also fixed and subjected to immunofluorescence microscopic analysis.

To explore the effects of serine proteases inhibitors, including nafamostat (Abcam), aprotinin (G-Biosciences), leupeptin (Bachem), and AEBSF (MP Biomedicals) on influenza infection, the inhibitor was delivered into the airway channel of influenza-infected chip (MOI = 0.1). Two days later, the virus samples were collected for detection of viral load and the vascular effluents were collected for analysis of cytokines and chemokines.

To test the efficacy of oseltamivir acid or antibodies directed against HA from H1N1 or H3N2, human Airway Chips infected with influenza virus (MOI = 0.1) were treated with 1 μM oseltamivir acid (Sigma-Aldrich), 10 μg/mL of anti-H1N1 HA antibody (Sino Biological), or 10 μg/mL of anti-H3N2 HA antibody (Abcam) under flow (60 μl/h) through the vascular channel. Two days later, virus samples were collected for detection of viral load, and cell layers were fixed and subjected to immunofluorescence microscopic analysis.

In the therapeutic window detection experiment, oseltamivir acid (1 μM), nafamostat (10 μM), or both were added to the influenza H1N1-infected Airway Chips (MOI = 0.1) at indicated times. Oseltamivir was perfused through the vascular channel, while nafamostat was introduced in 20 uL of PBS and incubated in the airway channel for 48 hours. Fluids samples were then collected from both channels for detection of viral load.

**Analysis of virus hemagglutinin (HA)**

A small volume (20 uL) of PBS containing 10 μM nafamostat, aprotinin, leupeptin, or AEBSF was introduced into the airway channels of Airway Chips when they were infected with influenza A/WSN/33 (H1N1) virus (MOI = 0.1). After 48 h, the airway epithelium and endothelium were harvested by perfusing 50 uL of RIPA Lysis buffer (Thermo Scientific) through the both channels, and the collected proteins were subjected to Western blot analysis using anti-H1N1 HA antibody.

For analysis of HA cleavage by serine proteases in presence/absence of nafamostat, MDCK cells ($5 \times 10^5$ cells per well in 6-well plates) were transfected with 2.5 μg serine protease expression plasmid or empty vector using TransIT-X2 Dynamic Delivery System (Mirus). One day later, the cells were infected with influenza A/WSN/33 (H1N1) virus (MOI = 0.01) in DMEM supplemented with 1% FBS and further cultured in presence or absence of 10 μM nafamostat. Two days post-infection, the supernatant was harvested and subjected to Western blot analysis using anti-HA1 antibody.

**Enzyme-linked immunosorbent assays (ELISAs)**

To detect the expression levels of serine proteases, including TMPRSS2, TMPRSS4, TMPRSS11D (HAT), TMPRSS11E (DESC1), total proteins were extracted from cultures of chips in RIPA Lysis buffer and analyzed using commercial ELISA kits for human TMPRSS2 (G-Biosciences), TMPRSS4 (G-Biosciences), TMPRSS11D (MyBioSource), and TMPRSS11E (MyBioSource).

**Analysis of cytokines and chemokines**

Vascular effluents from Airway Chips were collected and analyzed for a panel of cytokines and chemokines using custom ProcartaPlex assay kits (Invitrogen). Analyte concentrations were determined using a Luminex100/200 Flexmap3D instrument coupled with Luminex XPONENT software (Luminex, USA).

**Identification of drug-resistant virus strains**

Airway Chips were infected with amantadine- and oseltamivir-sensitive influenza A/WSN/33 virus (MOI = 0.01) and treated with 1 µM amantadine (Sigma-Aldrich), 1 µM oseltamivir acid, 10 µM nafamostat, or left untreated for 48 h. Amantadine or oseltamivir acid was perfused through the vascular channel under flow (60 µl/h), while nafamostat was diluted in 20 µL PBS and delivered into the airway channel. The progeny viruses were isolated by incubating the airway channel with 50 µL PBS for 1 h at 37℃,collecting the fluid, and then using it for infection in a new Airway Chip. This procedure was repeated up to 30 times, and after each passage, progeny virus yields were quantified using the plaque formation assay; the virus yield of untreated Airway Chips was set as 100%. When the progeny virus pool became resistant to drug treatment, the viruses were isolated through plaque purification and gene sequenced to identify gene mutations.

**Generation and identification of influenza reassortants**

Airway Chips were co-infected with pandemic influenza A/Netherlands/602/2009 (H1N1) virus (MOI = 2) and seasonal influenza A/Panama/2007/99 (H3N2) virus (MOI = 2) for 48 h, and then the progeny virus pool was collected by washing the airway channel with 50 uL PBS. The progeny viruses were isolated through plaque purification and gene sequenced to investigate any reassortment that occurred in their genomes. To evaluate the effects of reassortment on viral replication competence, parental virus strains and isolated reassortants were used to infect Airway Chips (MOI = 0.1), and their replication titers were quantified using the plaque formation assay at 48 h later. To evaluate the effects of reassortment on the efficacy of anti-HA (H3N2) antibody, parental virus strains and isolated reassortants were used to infect human Airway Chip (MOI = 0.1) in presence or absence of the antibody (10 µg/mL), and their replication titers were measured using the plaque formation assay.

To study reassortment between wild-type and drug-resistant viruses in the Airway Chip, wild-type, oseltamivir-resistant, and amantadine-resistant influenza/A/WSN/33 (H1N1) virus strains (MOI = 1) were used to infect Airway Chip in presence of oseltamivir and amantadine (1 μM). Two days later, their progeny viruses were isolated by plaque purification and gene sequenced.

**Statistical analysis**

Tests for statistically significant differences between groups were performed using a two-tailed unpaired Student's t-test. Differences were considered significant when the *P* value was less than 0.05 (*, $P<0.05$; **, $P<0.01$; ***, $P<0.001$; n.s., not significant). All results are expressed as means ± standard deviation (SD); error bars indicated $N>3$.

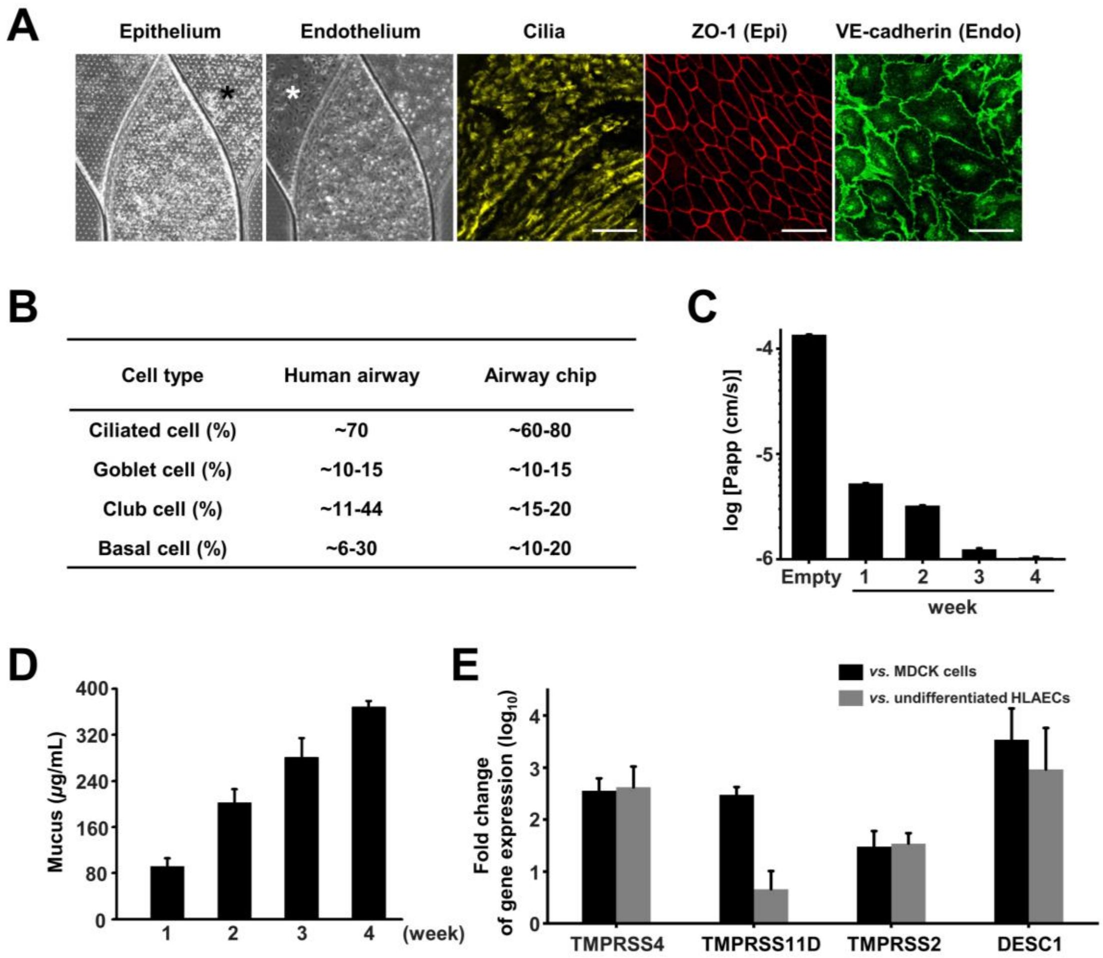


**Fig. S1. Characterization of human Airway Chip.** (**A**) Differentiation of human airway in chip. The epithelium exhibits well-structured cilia containing α–tubulin (yellow) and continuous ZO1-containing tight junctions (red), while the underlying endothelium contains continuous VE-cadherin-containing adherin junctions (green) linking adjacent cells (Bar, 50 μm). (**B**) Comparison of cell types between living human airway (*5*) and the human Airway Chip. (**C**) Barrier permeability (log $P_{app}$) of the Airway Chip assessed by the Cascade blue (607 Da) assay. (**D**) Mucus production at week 1, 2, 3, and 4 post-differentiation quantified using the Alcian Blue assay. (**E**) Fold changes in gene expression levels for various serine proteases in the well-differentiated human Airway Chip versus MDCK cells (one of the most commonly used cell lines in influenza studies) or undifferentiated HLAECs.

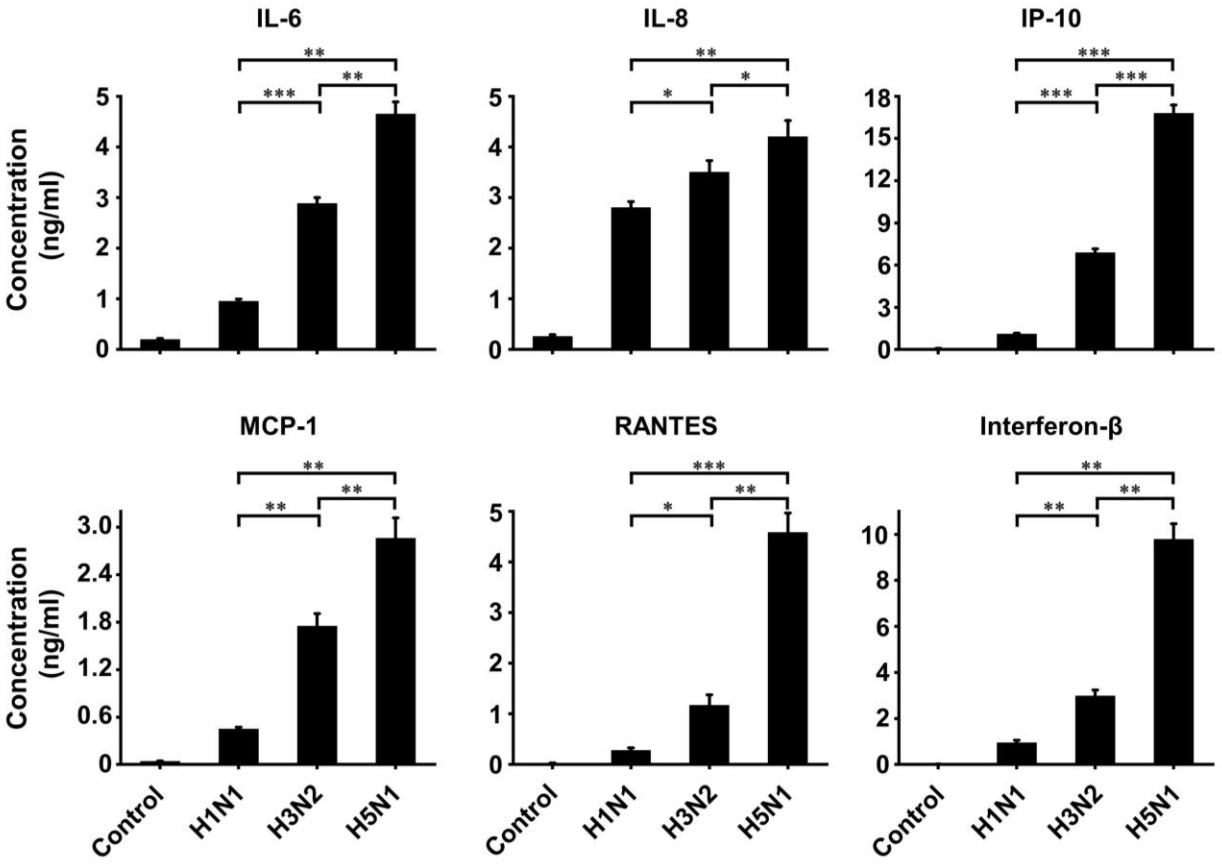


**Fig. S2. Production of cytokines and chemokines by the human Airway Chip at 48 h post-infection with different influenza virus strains, including H1N1, H3N2 and H5N1 (MOI = 0.1).** *, P<0.05; **, P<0.01; ***, P<0.001.

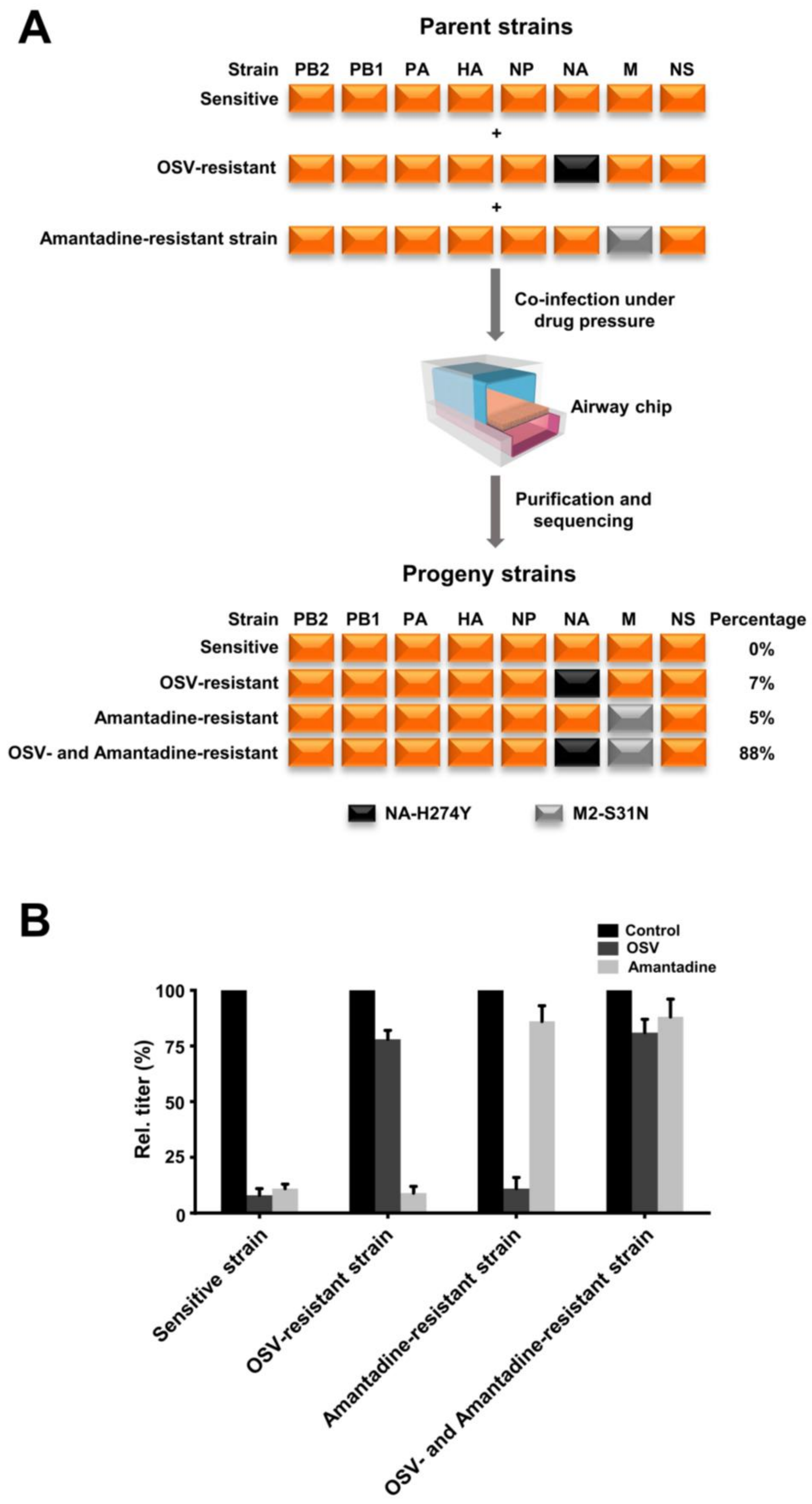

A
Parent strains
Strain PB2 PB1 PA HA NP NA M NS
Sensitive
+
OSV-resistant
+
Amantadine-resistant strain
Co-infection under drug pressure
Airway chip
Purification and sequencing
Progeny strains
Strain PB2 PB1 PA HA NP NA M NS Percentage
Sensitive 0%
OSV-resistant 7%
Amantadine-resistant 5%
OSV- and Amantadine-resistant 88%
NA-H274Y
M2-S31N
B
Control
OSV
Amantadine
Rel. titer (%)
100
75
50
25
0
Sensitive strain
OSV-resistant strain
Amantadine-resistant strain
OSV- and Amantadine-resistant strain

**Fig. S3. Generation of an influenza virus stain that is double resistant to both oseltamivir- and amantadine through reassortment in the Airway Chip.** (**A**) When drug-sensitive, OSV-resistant, and amantadine-resistant H1N1 influenza viruses (MOI = 1.0) were used to co-infect the same human Airway Chip in the presence of OSV and amantadine (1 μM), sequencing analysis of 100 progeny viruses isolated from the co-infected chips revealed that all progeny exhibited drug resistance and 3 genotypes were identified (%, relative incidence). Orange box indicates segment from the drug-sensitive virus strain; black box indicates segment from OSV-resistant strain; grey box indicates segment from amantadine-resistant virus strain. (**B**) Virus titer detection showing activity of OSV and amantadine (1 μM) against parental virus strain and the progeny reassortants.

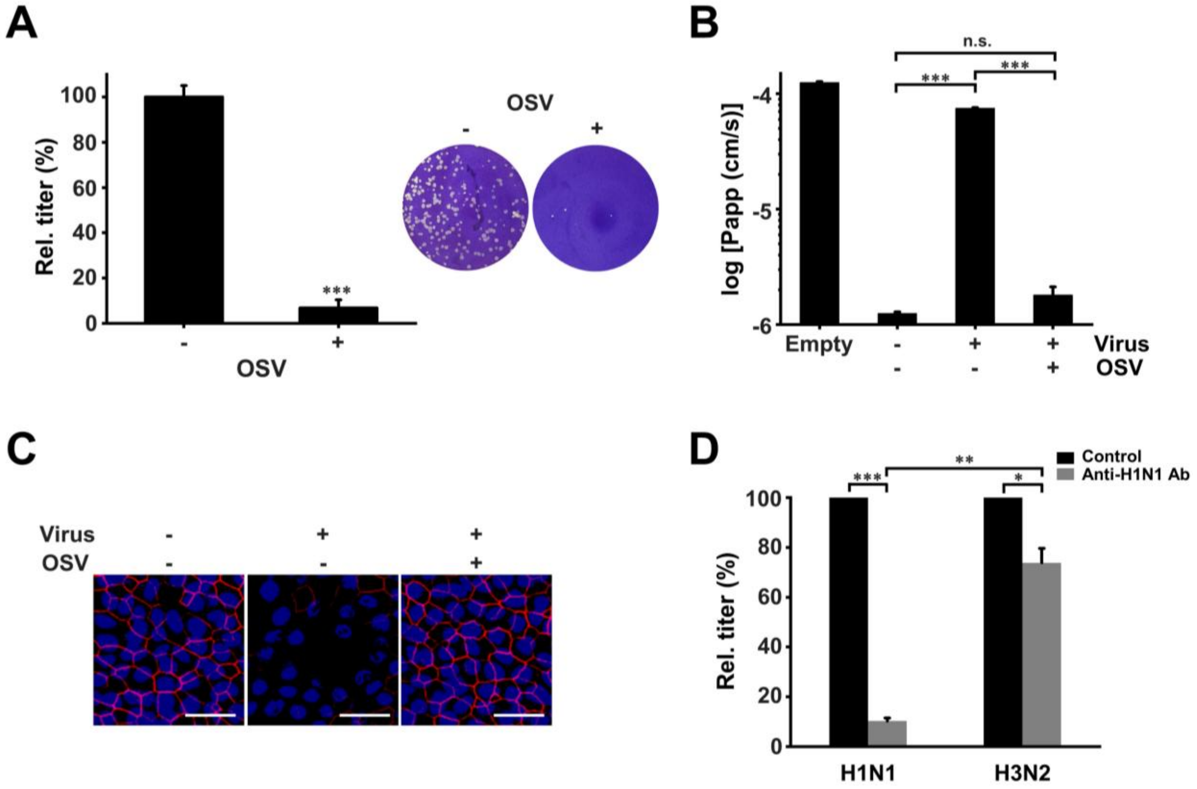


**Fig. S4. Inhibition of H1N1 influenza virus propagation in the human Airway Chip by oseltamivir and anti-H1N1 HA antibody.** (**A**) Graph (left) and photographs (right) showing plaque titers of progeny virus in the presence/absence of 1 μM oseltamivir acid (OSV). (**B**) Barrier permeability (log $P_{app}$) within the human Airway Chip measured 48 h post-infection with H1N1 (MOI = 0.1) with or without OSV. (**C**) Immunofluorescence micrographs showing the effects of OSV on ZO1-containing tight junctions in airway epithelium infected with H1N1 (MOI = 0.1) 48 h post-infection versus untreated. (**D**) Effects of 10 μg/mL of anti-H1N1 HA antibody on viral titer in Airway Chips infected with H1N1 and H3N2 virus. Bar, 50 μm. *, P<0.05; **, P<0.01; ***, P<0.001; n.s., not significant.

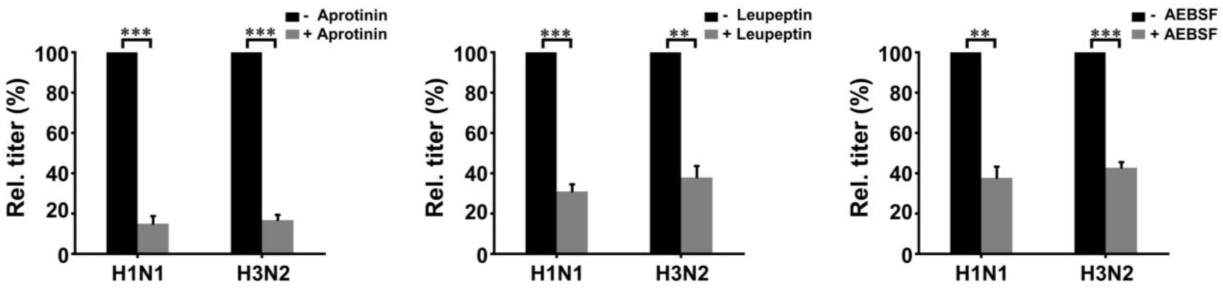


**Fig. S5. Aprotinin, leupeptin, and AEBSF suppress propagation of both H1N1 and H3N2 influenza viruses in human Airway Chip**. All drugs were added at 10 μM and viral titers were quantified 48 h post-infection. *, P<0.05; **, P<0.01; ***, P<0.001; n.s., not significant.

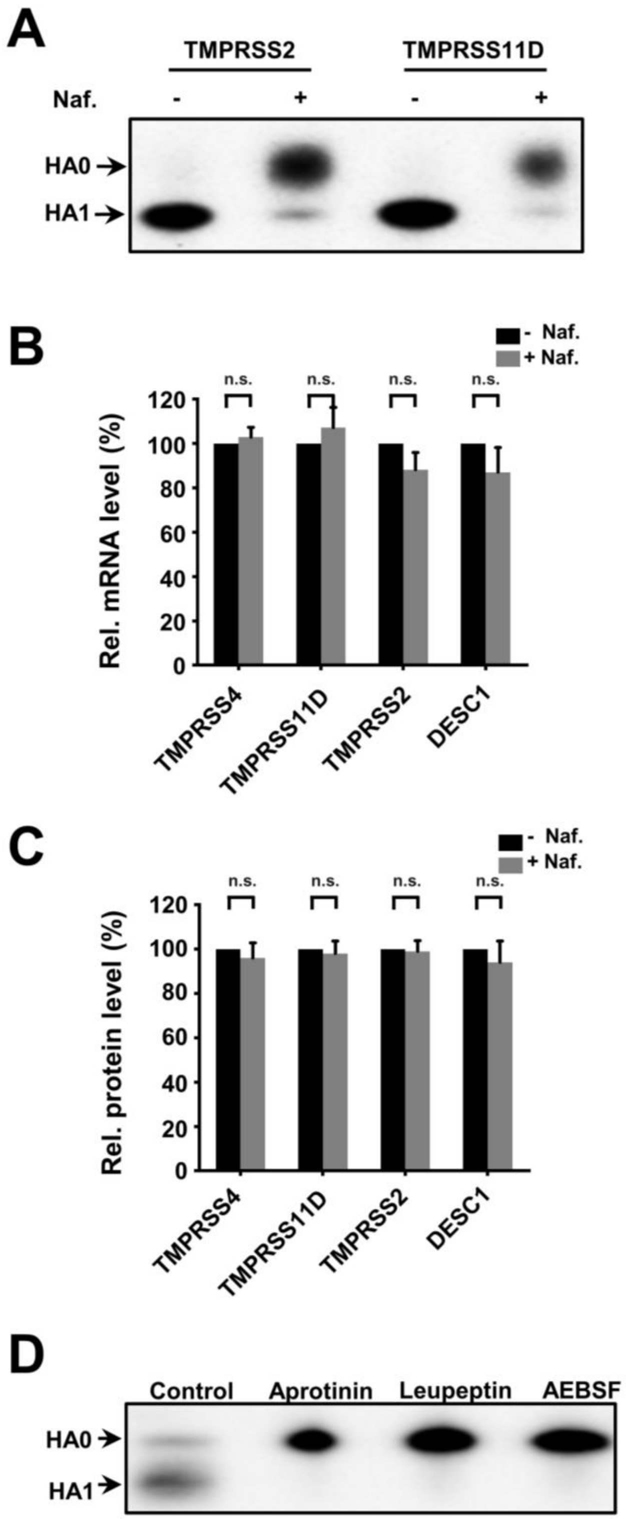


**Fig. S6. Exploration of the mechanism by which serine protease inhibitors suppress influenza virus infection.** (**A**) Western blots showing inhibition of TMPRSS2- and

TMPRSS11D-mediated cleavage of influenza virus HA0 by nafamostat (10 µM). (**B**) Effects of nafamostat (10 µM) on mRNA levels of the serine proteases, TMPRSS4, TMPRSS11D, TPMRSS2, and DESC1, in the human Airway Chip relative to untreated chips, as detected using qRT-PCR. (**C**) Effects of nafamostat (10 µM) on protein expression levels for the various serine proteases compared to untreated chips as detected using ELISA. (**D**) Western blot showing inhibitory effects of aprotinin, leupeptin, and AEBSF on the cleavage of influenza virus HA0 into HA1 and HA2 subunits in Airway Chip. n.s., not significant.

**Table S1. Antibodies used in this study.**

| Protein/Structure/Cell | Antibody | Vendor and Catalog |
|---|---|---|
| Tight junction | Alexa Fluor 594 anti-ZO-1 | Life Technologies, Cat# 339194 |
| Cilia | Alexa Fluor 647 anti-Acetyl-α-Tubulin | Cell Signaling Technology, Cat#81502 |
| VE-Cadherin | FITC anti-human VE-cadherin | BD Biosciences, Cat# 560411 |
| Goblet cell | Anti-Mucin5AC | Santa Cruz Biotechnology, Cat# sc-21701 |
| Club cell | Anti-human Uteroglobin/cc-10 | R&D Systems, Cat# MAB4218SP |
| Basal cell | Anti-Cytokeratin 5 | Sigma-Aldrich, Cat# SAB5300265 |
| Influenza NP | Anti-influenza NP | Invitrogen, Cat# MA516291 |
| ECM/Collagen | Anti-Collagen IV α1 | Novus Biologicals, Cat# NBP1-97716G |
| Neutrophil | Alexa Fluor 594 anti-human CD45 | Biolegend, Cat# 368520 |
| Influenza HA1 (H1N1) | Rabbit anti-influenza A H1N1 HA1 antibody | Sino Biological, Cat# 11692-T62 |
| Influenza HA (H3N2) | Mouse monoclonal [AT1B7] to Influenza A H3N2 HA antibody | Abcam, Cat# ab139361 |
| TMPRSS2 | Mouse anti-TMPRSS2 antibody | Novus Biologicals, Cat# H00007113-B01P |
| TMPRSS4 | Rabbit anti-TMPRSS4 antibody | Novus Biologicals, Cat# NBP1-56991 |
| TMPRSS11D | Mouse anti-TMPRSS11D antibody | Abnova, Cat# H00009407-B01 |
| TMPRSS11E | Rabbit anti-TMPRSS11E (DESC1) antibody | OriGene Technologies, Cat# TA350522 |
| Secondary antibody | Goat anti-mouse IgG, Alexa Fluor 488/594/647 | Life Technologies |
| | Goat anti-rabbit IgG, Alexa Fluor 488/594/647 | Life Technologies |
| | Goat anti-rabbit IgG H&L (HRP) | Abcam |
| | Goat anti-mouse IgG H&L (HRP) | Abcam |

**Supplementary Table 2. Primer sequences used for RT-qPCR analysis.**

| Gene | Primer | Sequence (5'-3') |
|---|---|---|
| TMPRSS2 | Forward | CTTTGAACTCAGGGTCACCA |
| | Reverse | TAGTACTGAGCCGGATGCAC |
| TMPRSS4 | Forward | TGCTTCAGGAAACATACCGA |
| | Reverse | CTGGAGTGAGCTCCTCATCA |
| TMPRSS11D | Forward | TACACAGGAATACAGGACTT |
| | Reverse | CTCACACCACTACCATCT |
| DESC1 | Forward | GTTGGTGGGACAGAAGTAGAAG |
| | Reverse | TGTAGGGAACAGGGCTAGAA |
| GAPDH | Forward | GAAGGTGAAGGTCGGAGTC |
| | Reverse | GAAGATGGTGATGGGATTTC |

**Movie S1. Real-time imaging of infection of the human Airway Chip by GFP-labeled influenza A/PR/8/34 (H1N1) virus.** The human Airway Chip was inoculated with GFP-labeled PR8 virus (MOI = 0.01) and cultured for 36 h; virus infection is indicated by the progressive increase in GFP-positive cells. (**A**) GFP signal, (**B**) merge of GFP and bright field.
